# Gene regulatory architecture in antibiotic resistance is optimized for specific dynamic regimens

**DOI:** 10.64898/2026.11.03.764017

**Authors:** Hao Geng, Daniel Schultz

## Abstract

Cellular responses to antibiotics face a fundamental trade-off: resistance must be deployed quickly enough to prevent the accumulation of drug inside the cell, but expression of resistance genes is costly in the absence of the drug. Therefore, antibiotic responses are typically regulated, controlling not only whether resistance is expressed, but also when, how strongly, and for how long. These regulatory strategies are implemented through the strength and arrangement of promoters and operators. However, it is still not clear how the dynamics of drug exposure select for specific regulatory strategies. Here, we develop a dynamical model of a typical antibiotic response, where a transcriptional repressor senses the drug and regulates expression of a resistance protein, coupled to the effects of the drug on cell growth and gene expression. We define a cost function that weighs the different burdens imposed on the cell over the course of the response, and perform *in silico* optimization of the regulatory parameters under different selective pressures for three architectures of increasing complexity: constitutive expression of resistance, repression by a constitutively expressed repressor, and repression by a self-regulated repressor. We find that strong drug exposures favor constitutive expression of resistance, high costs of basal expression favor repression by a constitutive repressor, and self-regulation of the repressor is favored only when the response must provide protection while also restraining overexpression of resistance. Stochastic simulations further show that self-regulation of the repressor protects cells against fluctuations in repressor levels, which cause unnecessary induction of resistance before exposure and failed induction after exposure. This framework links the dynamics of the environment to the evolution of gene regulation, providing a basis to predict which regulatory strategies are expected to arise under specific selective pressures.

**Significance:** Bacteria regulate the expression of antibiotic resistance genes to avoid the costs of producing resistance when it is not needed. However, regulated responses take time to activate, which can be fatal when cells are suddenly exposed to large drug doses. Here, we use a mathematical model of antibiotic responses to identify which regulatory strategies are optimal under different selective pressures. We show that the choice between constitutive and regulated expression of resistance, and the use of self-regulation by the repressor, depend on the dynamics of drug exposure and on which costs dominate the response. Noise in gene expression further favors self-regulation of the repressor near the limit of tolerable drug exposures. These results help explain the diversity of regulatory architectures found in natural resistance mechanisms, and the frequent loss of regulation in pathogens exposed to antibiotic treatments.

## Introduction

Microbes are equipped with regulated mechanisms that sense changes in their environment and induce the appropriate cellular responses^1–3^. In the case of antibiotic responses, resistance genes are typically repressed in the absence of the drug, avoiding the costs associated with their expression^4–6^. Therefore, when cells are suddenly exposed to antibiotics, they have only a short window of time to induce resistance before drug accumulation impairs growth and gene expression^7,8^. However, an optimal response does not depend solely on whether the drug is eventually removed from the cell, but by its full time course, which includes how much resistance is expressed before, during, and after exposure, and how well growth is maintained and restored as the drug accumulates and is subsequently cleared^9–11^.

This problem of designing optimized regulation of antibiotic responses involves fundamental trade-offs. Constitutive expression of resistance provides immediate protection upon drug exposure, but imposes a continuous burden on the cell in the absence of the drug^4,12^. Transcriptional repression reduces this burden by keeping resistance low before exposure, but delays activation of the response, which can impair growth and slow recovery^7,13^. The relative advantage of each strategy depends on which costs are most critical in a given environment^14,15^. Optimality principles have been used to explain the expression levels of individual proteins^9,16^ and the choice between different modes of gene regulation^17–20^, and dynamic optimization has been used to predict the regulation of metabolic pathways^21–24^. However, we still lack a framework to understand how the demands imposed by the full time course of a response shape the architecture of its regulation^25^: whether protection should be present before exposure, whether expression should remain repressed until needed, and whether additional control of the transcription factor itself is advantageous.

In previous work, we developed models of antibiotic responses that link the expression of resistance genes to the effects of the drug on cell growth and metabolism^8^, and used optimization to show how different selective pressures favor different regulatory parameters within a fixed regulatory architecture^26^. We also showed experimentally that the dynamics of drug administration dictate the evolution of antibiotic responses, often leading to qualitatively different forms of regulation^13^. Here, we combine these approaches to determine how environmental demands shape the architecture of regulation itself. We consider a general antibiotic response consisting of a drug, a resistance protein that processes the drug, and a transcriptional repressor that senses the drug and represses expression of resistance (**Fig. 1A**), modeled after the tetracycline resistance *tet* operon^27–31^. We focus on sudden increases in extracellular drug concentration, which place particular emphasis on the timing of activation and on the ability of the cell to maintain and recover growth. We translate the selective pressures acting on the response into a cost function that weighs key metrics of the response, and identify the regulatory parameters that minimize this cost for three architectures of increasing complexity (**Fig. 1B**). Finally, we use stochastic simulations to determine how noise in gene expression affects the choice of regulatory architecture.

**Figure 1.**
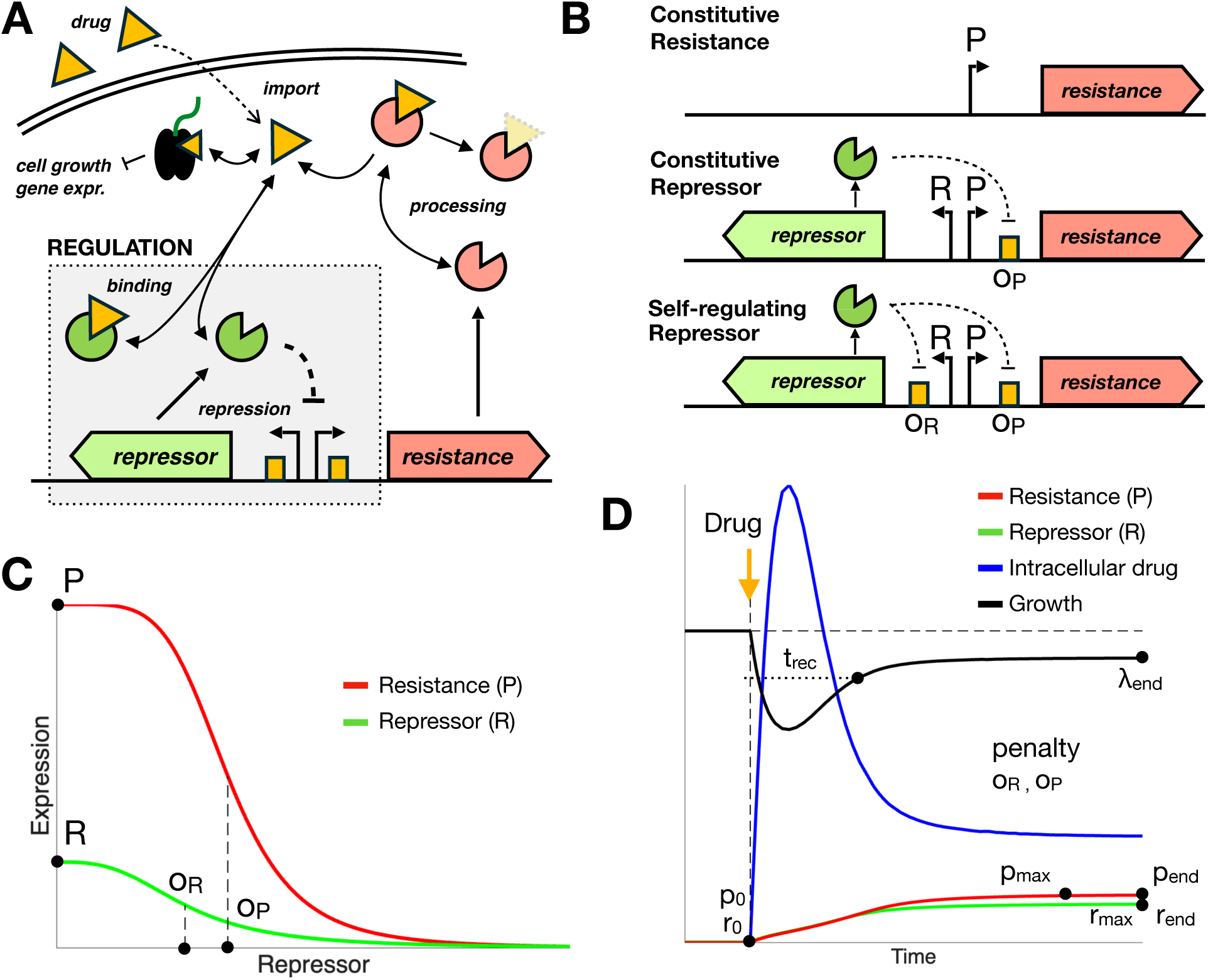
Model of a regulated antibiotic response. (**A**) Drug is imported into the cell, where it binds the repressor, which loses affinity for its binding sites and releases expression of resistance. The resistance protein processes the drug. Free repressor represses expression of resistance and, in the self-regulating architecture, of itself. The drug also inhibits growth and gene expression (not shown). (**B**) Regulatory architectures considered in this study. *Constitutive resistance*: resistance is expressed from an unregulated promoter. *Constitutive repressor*: a constitutively expressed repressor represses resistance through operator *O*_P_. *Self-regulating repressor*: the repressor also represses its own promoter through operator *O*_R_. (**C**) Expression rates of resistance (red) and repressor (green) as functions of free repressor, described by Hill functions. Promoter strengths *P* and *R* set the maximal expression rates, and operator thresholds *O*_P_ and *O*_R_ set the concentrations of free repressor resulting in half-maximal repression. (**D**) Typical time course of a response to a sudden drug exposure at time zero (orange arrow), showing resistance (red), repressor (green), intracellular drug (blue), and growth (black). The metrics used in the cost function are indicated: basal (*p*_0_, *r*_0_), maximal (*p*_max_, *r*_max_), and final (*p*_end_, *r*_end_) levels of resistance and repressor, final growth rate *λ*_end_, recovery time *t*_rec_, and the penalty on operator strength.

## Results

### Model of antibiotic response coupled to cell growth

We model the response by a system of three differential equations describing the concentrations of intracellular drug *d*, resistance protein *p*, and repressor *r* (**Fig. 1A**, **Methods**)^8,26^. The drug enters the cell at a rate proportional to the difference between extracellular and intracellular concentrations, is processed by the resistance protein following Michaelis–Menten kinetics, and is diluted by cell growth. Inside the cell, the drug binds the repressor, which loses affinity for its binding sites in the DNA. Since binding and unbinding of the drug happen much faster than the other processes, we consider the free forms of drug and repressor to be in chemical equilibrium with the bound complex. Free repressor regulates the expression of resistance, and possibly of itself, according to Hill functions with Hill coefficient *n* = 2, reflecting the binding of repressor dimers (**Fig. 1C**). Each regulated gene is then described by two parameters: a promoter strength that sets its maximal expression rate (*P* for resistance and *R* for the repressor), and an operator threshold, the concentration of free repressor that results in half-maximal repression (*O*_P_ and *O*_R_, respectively), which reflects the affinity between the operator and the repressor.

Following exposure, the drug also inhibits translation, reducing cell growth and gene expression. We incorporate these effects using the proteome partition framework of Scott *et al.*^32,33,8^. The growth rate *λ* depends on the translational capacity of the cell, which decreases with the concentration of free intracellular drug, and expression of resistance and repressor is modulated by a factor *f* = *λ*^2^ that accounts for both the reduction in growth and the reallocation of the proteome towards ribosomal proteins ^34,35^. This coupling creates a positive feedback: failure to quickly deploy resistance results in higher drug accumulation, which further reduces growth and expression of resistance. At high drug concentrations, this feedback results in bistability between recovered and arrested states^7,8,36^. Additional model parameters are based on the tetracycline response in *E. coli*, and all variables are rescaled so that time is measured in units of the inverse of the drug-free growth rate (**Methods**, **Supplementary Text S1**, **Tables S1–S3**).

A typical response to a sudden increase in extracellular drug concentration proceeds as follows (**Fig. 1D**). Before exposure, resistance is kept at a low basal level *p*_0_ by the repressor, which is at its basal level *r*_0_. Upon exposure, the drug enters the cell and binds the repressor, releasing expression of resistance. Intracellular drug accumulates and growth decreases until enough resistance protein is expressed to reduce drug levels. Growth then recovers, repression is progressively restored, and the levels of all components relax to new steady states in the presence of the drug. We consider three regulatory architectures of increasing complexity (**Fig. 1B**): (1) *constitutive resistance*, where resistance is expressed at a constant rate without repression; (2) *constitutive repressor*, where resistance is repressed by a repressor that is expressed at a constant rate; and (3) *self-regulating repressor*, where the repressor also represses its own expression, as in the *tet* operon^29,37,38^.

### Selective pressures translated into a cost function

Different environments place different demands on an antibiotic response, depending on how often, how suddenly, and at what concentrations the drug is encountered. We therefore represent each environment as a combination of selective pressures that reflects the dynamics of drug exposure, with each pressure acting on a different phase of the response. In the absence of the drug, basal expression of resistance imposes a burden on the cell. Upon exposure, transient high levels of resistance and repressor can also be costly^4,12^, while delayed recovery and reduced steady-state growth are harmful long-term consequences of drug toxicity. To represent these pressures, we extract a set of metrics from the time course of the response (**Fig. 1D**, **Table S4**): basal (*p*_0_, *r*_0_), maximal (*p*_max_, *r*_max_), and final (*p*_end_, *r*_end_) levels of resistance and repressor, the final growth rate *λ*_end_, and the recovery time *t*_rec_, measured from the growth minimum until growth recovers to 80% of its final level. We also include a penalty on operator strength (1/ *O*_P_ and 1/*O*_R_), reflecting the evolutionary cost of maintaining tight binding sites and preventing the optimization from defaulting to excessively strong repression. The cost *C* of a response is then a weighted sum of these metrics (**Methods**). For simplicity, we assume that the cost grows linearly with each metric, since any cost that increases monotonically with every metric has its optima on the same Pareto front^26^. The relative weights reflect the selective pressures acting in each environment. For instance, when drug exposures are rare, cells spend most of their time in the absence of the drug, and a high weight on basal resistance *p*_0_ reflects the pressure to keep resistance repressed by default. Responses that fail to recover growth (*λ*_end_ < 0.2) or that do not stabilize within the simulated time are considered unviable. For each architecture, drug concentration, and set of weights, we then optimize the regulatory parameters (*P*, *R*, *O*_P_, *O*_R_) to minimize *C*, using a global search algorithm followed by local refinement (**Methods**, **Supplementary Text S3**).

### Constitutive expression of resistance reveals trade-off between drug toxicity and cost of resistance

We begin with the simplest architecture, where resistance is expressed constitutively and the only regulatory parameter is the promoter strength *P* (**Fig. 2A**). In this case, the basal, maximal, and final levels of resistance cannot be tuned independently, and optimization reduces to the choice of a single expression level that balances the cost of resistance against the protection it provides. We therefore consider a cost composed only of the basal level of resistance and the final growth rate, *C* = *w*_p0_*p*_0_ + *w*_λ_/*λ*_end_, and vary the relative weight on resistance *w*_p0_/*w*_λ_ together with the extracellular drug concentration *D* (**Fig. 2B–D**). As the drug concentration increases, the optimal expression of resistance increases, reflecting the increasing benefit of protection (**Fig. 2D**). Increasing the cost of resistance shifts the optimum towards lower expression, at the expense of lower growth (**Fig. 2C**).

**Figure 2.**
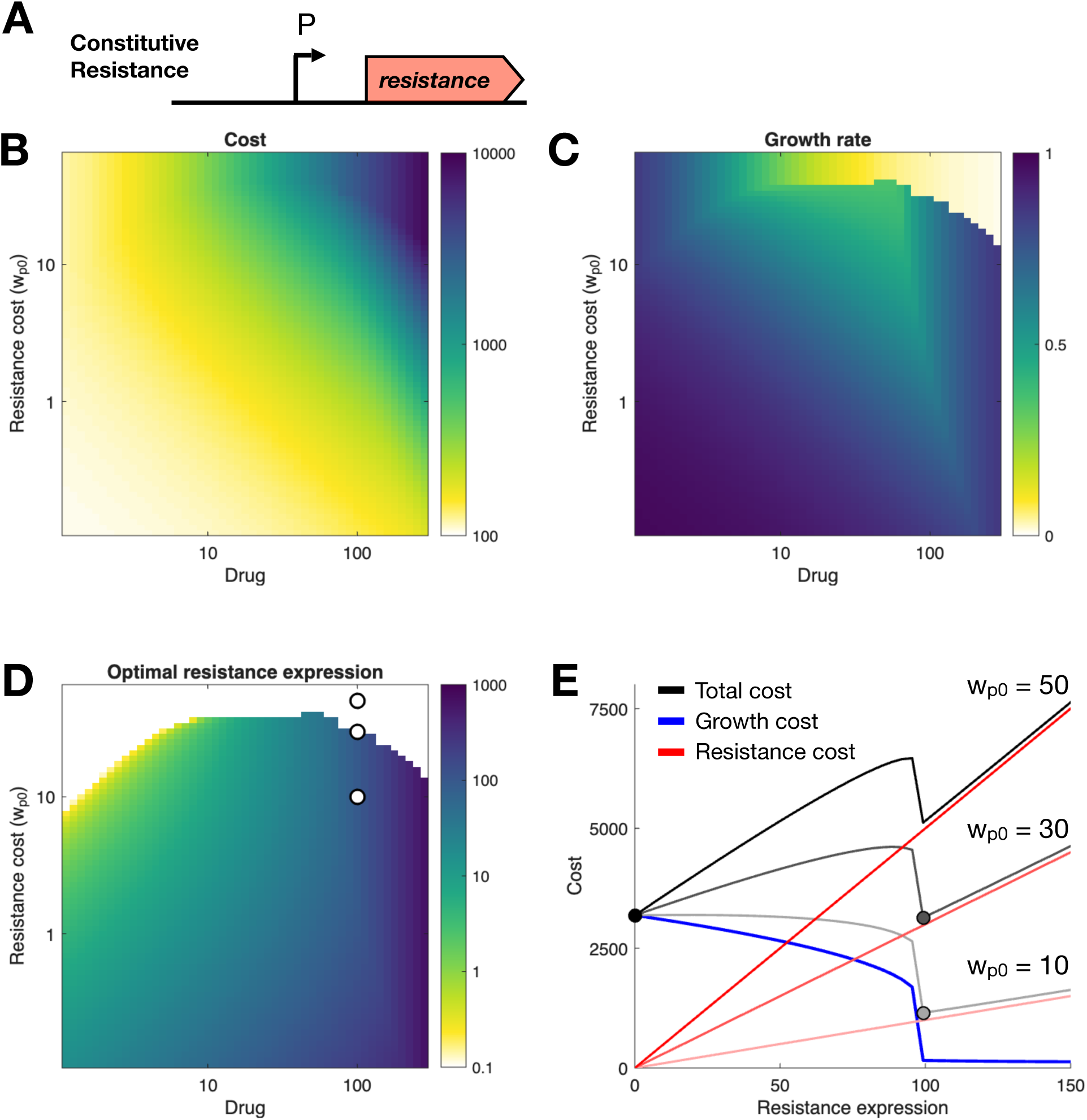
Optimal constitutive expression of resistance. (**A**) Constitutive resistance architecture, with promoter strength *P*. (**B–D**) Total cost (**B**), final growth rate (**C**), and optimal expression of resistance (**D**) as functions of the extracellular drug concentration *D* and of the weight on basal resistance *w*_p0_, for a cost *C* = *w*_p0_*p*_0_ + 100/*λ*_end_. (**E**) Components of the cost as functions of the level of resistance at *D* = 100, for *w*_p0_ = 10, 30, and 50 (circles in **D**): cost of expressing resistance (red lines; slope increases with *w*_p0_), growth cost 100/*λ*_end_ (blue), and total cost (black and gray). Circles mark the optimal levels of resistance. Above a threshold cost of resistance, the optimum switches from the minimum level of resistance that allows recovery of growth to no expression of resistance.

However, the optimal expression of resistance does not decrease smoothly with its cost. Above a threshold cost, the optimal strategy abruptly switches to no expression of resistance, with the cell accepting low final growth (white region at the top of **Fig. 2D**). To understand this transition, we examined how each component of the cost varies with the level of resistance at a fixed drug concentration (**Fig. 2E**). Because of the positive feedback between drug accumulation and growth, final growth is close to zero for low levels of resistance, and increases sharply once resistance is sufficient to escape arrest. The total cost therefore has two local minima: one at zero expression, where the cell is arrested, and one just above the level of resistance needed for recovery. As the cost of resistance increases, the cost of the resistant solution increases linearly until it exceeds the cost of arrest, and the optimum switches to the minimum corresponding to no resistance. As a result, the range of drug concentrations over which growth can be maintained becomes narrower as the cost of resistance increases (**Fig. 2C**). Therefore, constitutive expression of resistance cannot separate protection during exposure from its cost in the absence of the drug, and growth collapses beyond a critical drug concentration that depends on the cost of resistance.

### Repression by a constitutive repressor is favored when basal expression of resistance is costly

We next asked under what conditions it becomes advantageous to regulate the expression of resistance (**Fig. 3**). We introduced a constitutively expressed repressor that represses resistance (**Fig. 3A**), and optimized the promoter strengths of resistance and repressor (*P*, *R*) and the operator threshold (*O*_P_). Since constitutive resistance is a limiting case of this architecture, the optimization can also discard repression altogether, since optimized architectures with low repressor level *R* and high operator threshold *O*_P_ are equivalent to constitutive resistance. In addition to the basal level of resistance and final growth, the cost now includes the maximal levels of resistance and repressor, the recovery time, and the operator penalty, and we vary the weight on basal resistance *w*_p0_ together with the drug concentration (**Methods**, **Table S5**). Increasing *w*_p0_ therefore represents environments where expression of resistance in the absence of the drug is especially costly, such as when drug exposures are rare.

**Figure 3.**
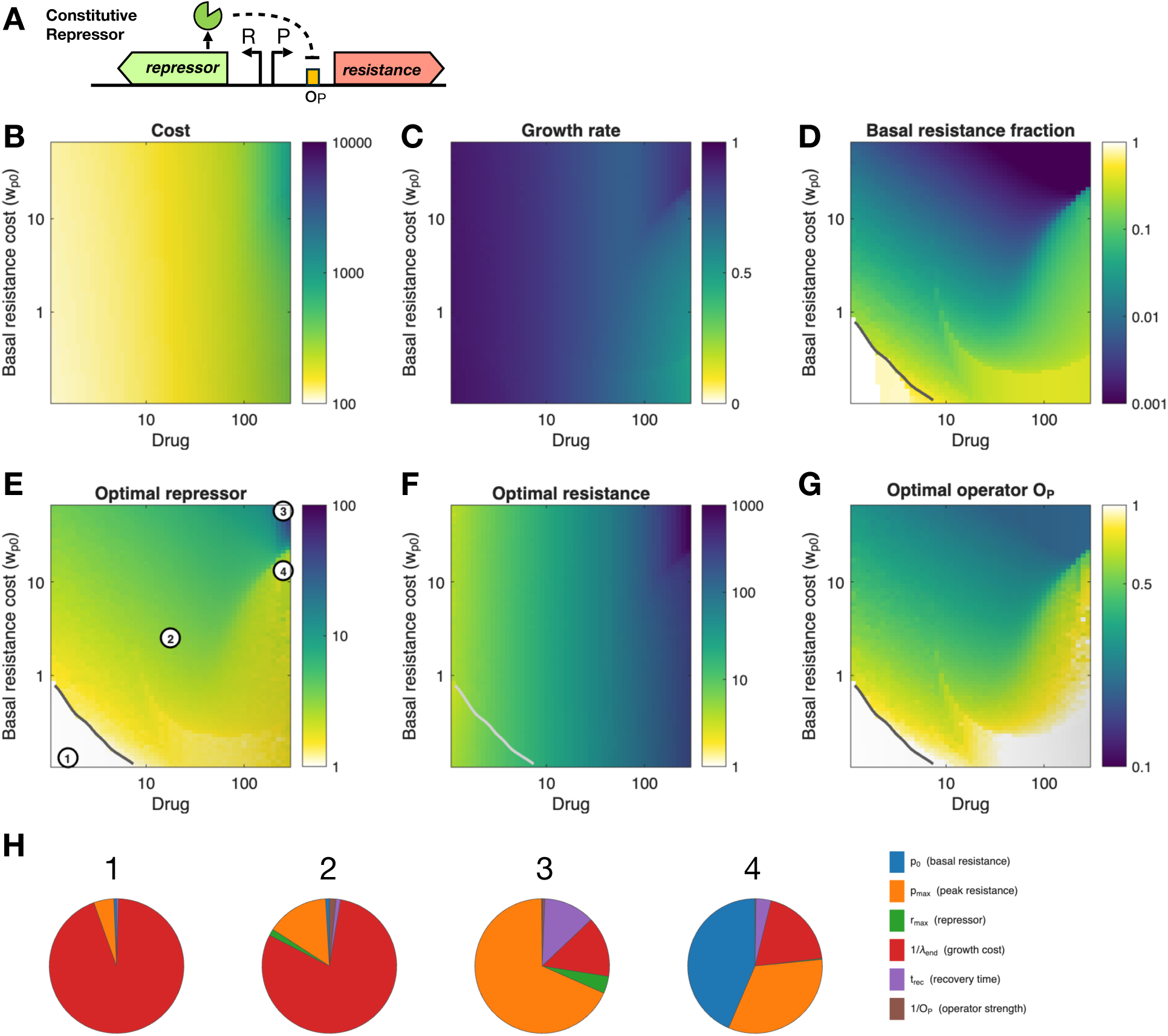
Optimal regulation by a constitutive repressor. (**A**) Constitutive repressor architecture, with promoter strengths *P* and *R* and operator threshold *O*_P_. (**B–G**) Total cost (**B**), final growth rate (**C**), basal fraction of resistance *p*_0_/*P* (**D**), and optimal promoter strength of the repressor *R* (**E**), promoter strength of resistance *P* (**F**), and operator threshold *O*_P_ (**G**), as functions of the drug concentration *D* and of the weight on basal resistance *w*_p0_, for a cost *C* = *w*_p0_*p*_0_ + *p*_max_ + *r*_max_ + 100/*λ*_end_ + *t*_rec_ + 1/*O*_P_. Gray lines mark the boundary below which the optimal response loses repression. (**H**) Breakdown of the cost into its components at the four points marked in **E**.

When basal expression of resistance is costly, the optimal response does not simply reduce the expression of resistance, but becomes increasingly inducible. As *w*_p0_ increases, the optimal repressor level increases (**Fig. 3E**) and the operator threshold decreases (**Fig. 3G**), lowering basal resistance to a small fraction of its maximal expression (**Fig. 3D**), while the promoter strength of resistance remains high (**Fig. 3F**). The result is a response that keeps resistance low before exposure, but can still strongly induce it once the drug accumulates in the cell. In contrast, when basal resistance is cheap and drug concentrations are low, there is little benefit in keeping resistance off, and the optimal response loses repression entirely (region below the gray lines in **Fig. 3D–G**).

This logic changes as the drug concentration increases. With higher drug doses, the benefit of low basal resistance is progressively outweighed by the cost of delayed activation, since the time required for the drug to accumulate and inactivate the repressor results in a larger growth burden. Accordingly, at high drug concentrations strong repression is only maintained when basal resistance is very costly, and the basal fraction of resistance otherwise approaches constitutive levels (**Fig. 3D**). The breakdown of the cost into its components shows how different burdens dominate in each regime (**Fig. 3H**). At low drug concentrations and low costs of resistance, the cost is dominated by growth (point 1), with peak expression of resistance becoming more important as the drug concentration increases (point 2). At high drug concentrations, the cost is dominated by peak resistance when basal resistance is very costly (point 3), while basal resistance itself becomes a major cost at intermediate weights (point 4), where the response is only partially repressed. Thus, repression by a constitutive repressor is favored when basal expression is costly and drug exposures are rare, since the benefit of keeping resistance off between exposures outweighs the cost of delayed induction. When exposures become too frequent or too severe, the cost of delayed induction dominates, and the optimal strategy shifts towards constitutive expression of resistance.

### Self-regulation of the repressor is favored only under demanding regimes

Finally, we considered the architecture where the repressor also represses its own expression (**Fig. 4A**). Here, we fixed the weight on basal resistance (*w*_p0_ = 10) and varied the weight on maximal resistance *w*_pmax_ together with the drug concentration, optimizing all four regulatory parameters (*P*, *R*, *O*_P_, *O*_R_). If self-regulation is not beneficial, the optimization drives the operator threshold of the repressor *O*_R_ to high values, and the architecture reduces to a constitutive repressor. Therefore, the region where self-regulation is used can be identified by the basal level of repressor relative to its maximal expression, *r*_0_/*R* (**Fig. 4D**).

**Figure 4.**
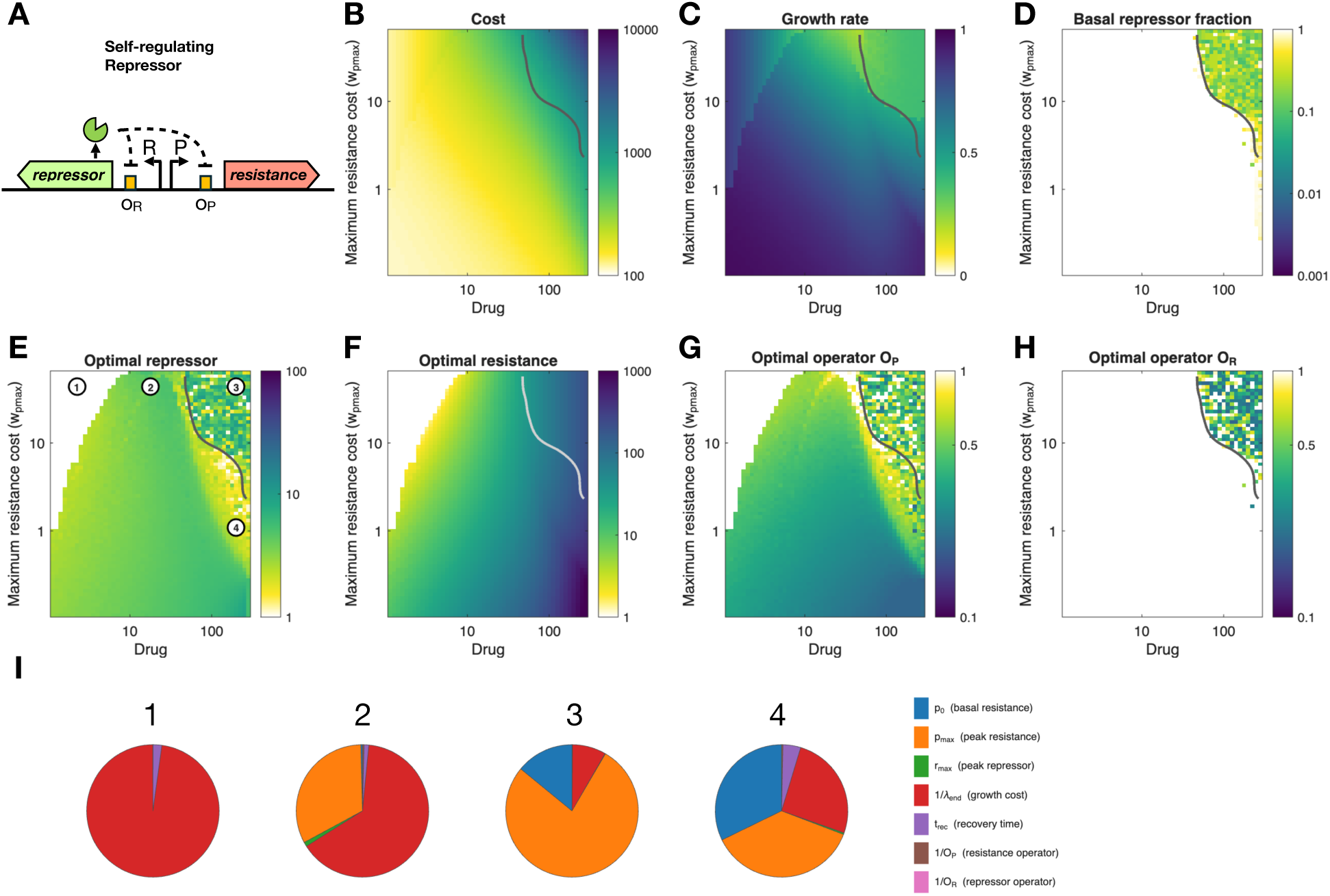
Optimal regulation by a self-regulating repressor. (**A**) Self-regulating repressor architecture, with promoter strengths *P* and *R* and operator thresholds *O*_P_ and *O*_R_. (**B–H**) Total cost (**B**), final growth rate (**C**), basal fraction of repressor *r*_0_/*R* (**D**), and optimal promoter strength of the repressor *R* (**E**), promoter strength of resistance *P* (**F**), operator threshold of resistance *O*_P_ (**G**), and operator threshold of the repressor *O*_R_ (**H**), as functions of the drug concentration *D* and of the weight on maximal resistance *w*_pmax_, for a cost *C* = 10*p*_0_ + *w*_pmax_*p*_max_ + *r*_max_ + 100/*λ*_end_ + *t*_rec_ + 1/*O*_P_ + 1/*O*_R_. Gray lines delimit the region where self-regulation of the repressor is used; outside this region, the optimal response uses a constitutive repressor (*r*_0_/*R* = 1, blank in **D** and **H**). (**I**) Breakdown of the cost into its components at the four points marked in **E**.

The resulting parameter maps show that self-regulation is only favored in a limited region of high drug concentrations and high costs of peak resistance (region above the gray line in **Fig. 4B–H**). Over most of the parameter space, the optimal response uses a constitutive repressor, with control concentrated on the resistance promoter and its operator (**Fig. 4E–G**). The promoter strength of the resistance protein increases with the drug concentration across the viable region, and depends only weakly on the cost of peak resistance (**Fig. 4F**). Therefore, the need for protection remains the main determinant of the capacity to express resistance, even when overexpression of resistance is penalized. Instead, higher costs of peak resistance are met by adjusting the operator threshold of resistance (**Fig. 4G**), which controls how much of this capacity is released during the response. At low drug concentrations and high costs of peak resistance, the benefit of protection does not justify expression of resistance, which is turned off (white region at the top left of **Fig. 4E–G**).

Self-regulation of the repressor is recruited only at high drug concentrations, when the cost of peak resistance is also high (**Fig. 4D, H**). In this regime, the response must simultaneously provide protection against a large drug dose, restrain overexpression of resistance, and preserve growth. The cost breakdown shows that peak resistance dominates the cost in this region (point 3, **Fig. 4I**), while growth dominates at lower drug concentrations (points 1 and 2), and basal resistance becomes a significant cost when peak resistance is cheaper (point 4). Within the self-regulated region, the optimal parameters vary considerably between neighboring points (**Fig. 4D–H**), indicating a nearly flat cost landscape, where different combinations of parameters result in similar costs. Therefore, self-regulation of the repressor is not a requirement for an efficient response, but provides a slight advantage only under the most demanding regimes.

Taken together, these results show that the optimal regulatory strategy changes systematically with the dynamical demands imposed by the environment. Optimization does not simply tune expression levels within a fixed circuit, but shifts the response between constitutive expression of resistance, repression by a constitutive repressor, and repression by a self-regulated repressor. Strong drug exposures favor pre-existing protection, and therefore constitutive expression of resistance. High costs of basal resistance favor repression by a constitutive repressor, which keeps resistance off between exposures. Additional costs of peak resistance favor the recruitment of self-regulation of the repressor, which helps restrain transient overexpression of resistance.

### Noise in repressor expression favors self-regulation of the repressor

The optimization above treats the response as deterministic. However, expression of both the repressor and the resistance protein is stochastic at the single-cell level^39,40^, and the low copy numbers of transcription factors makes expression of regulated genes particularly noisy^8,41,42^. These fluctuations are amplified during antibiotic exposure, where they can lead to the coexistence of recovered and arrested cells within isogenic populations^7,8^. We therefore asked whether noise changes the cost of each optimized architecture, and whether it can change which architecture is preferred.

We added noise to the synthesis of either the repressor or the resistance protein, using a chemical Langevin description in which the amplitude of fluctuations scales with the square root of the synthesis rate^43^, and is set by a relative factor *σ*_R_ for the repressor and *σ*_P_ for resistance (**Methods**, **Supplementary Text S4**). Noise is active for a period before drug exposure as well as during exposure, so the state of each cell when the drug arrives includes any spontaneous de-repression accumulated beforehand. For each architecture, drug concentration, and noise level, we simulated 1000 cells exposed to identical noise realizations, and averaged the cost function over all cells. Cells whose final growth fell below 20% of the drug-free rate (*λ*_end_ < 0.2) were counted as not surviving, and received the same low-growth penalty used in the deterministic optimization. We compared circuits optimized independently under the same cost function as in **Fig. 4**, at a weight on maximal resistance *w*_pmax_ = 0.5. Since optimization of the self-regulating architecture degenerates into the constitutive repressor when self-regulation is not beneficial, we enforced self-regulation by fixing the strength of repressor self-repression at *r*_0_/*R* = 0.2, the typical value within the region of **Fig. 4** where self-regulation is used, and optimized the remaining parameters. In the absence of noise, this self-regulated circuit costs slightly more than the constitutive-repressor circuit at all drug concentrations, from 6% more at low concentrations to less than 1% at the highest (**Table S6**). Therefore, without noise, the constitutive repressor is preferred everywhere along this range.

Noise in repressor expression changes this picture at high drug concentrations (**Fig. 5**). At intermediate drug concentrations (point 1, *D* = 50), the two architectures respond almost identically (**Fig. 5A–C**). Repressor noise produces small, transient bursts of resistance before exposure and some spread in the timing of induction, but neither effect substantially changes the cost, and the constitutive repressor remains 5–8% cheaper at all noise levels. As the drug concentration increases (point 2, *D* = 158), induction of resistance is delayed by a prolonged phase of growth arrest (**Fig. 5D–F**). Repressor noise then severely spreads the induction times of individual cells, and resistance at the time of exposure rises almost 10-fold above its noise-free basal level in the constitutive-repressor circuit, compared with 3-fold in the self-regulated circuit. As a result, the self-regulated architecture becomes 8–13% cheaper at strong repressor noise (*σ*_R_ ≥ 2). The difference is largest at the highest drug concentrations (point 3, *D* = 224; **Fig. 5G–I**).

**Figure 5.**
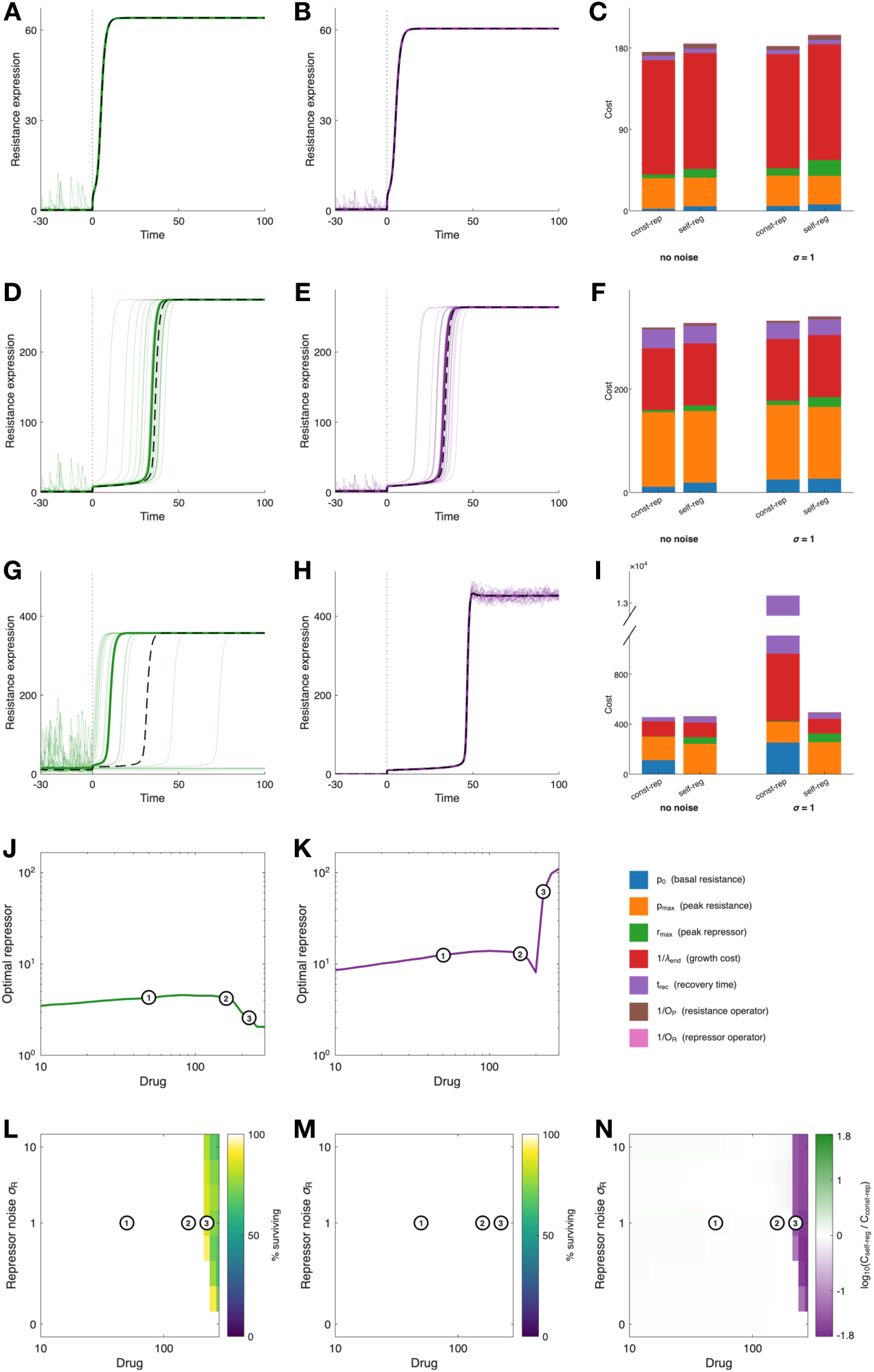
Noise in repressor expression favors self-regulation of the repressor. Circuits were optimized under the cost of Fig. 4 with *w*_pmax_ = 0.52, with the self-regulating repressor constrained to *r*_0_/*R* = 0.2. (**A–I**) Responses to a drug exposure at time zero for *D* = 50 (**A–C**, point 1), *D* = 158 (**D–F**, point 2), and *D* = 224 (**G–I**, point 3), for the constitutive repressor (**A, D, G**, green) and the self-regulating repressor (**B, E, H**, purple), with repressor noise *σ*_R_ = 1 starting 50 time units before exposure. Thin lines show expression of resistance in 20 single cells, thick lines the median of 1000 cells, and dashed lines the noise-free response. (**C, F, I**) Breakdown of the cost without noise and with *σ*_R_ = 1, averaged over 1000 cells, including cells that collapse. The broken axis in **I** shows the cost of the constitutive repressor, dominated by the penalty assigned to collapsed cells. (**J, K**) Optimal promoter strength of the repressor *R* for the constitutive (**J**) and self-regulating (**K**) repressor, which does not depend on noise. (**L, M**) Percentage of surviving cells (*λ*_end_ ≥ 0.2) for the constitutive (**L**) and self-regulating (**M**) repressor, as functions of the drug concentration *D* and repressor noise *σ*_R_. (**N**) Preferred architecture, measured by log_10_(*C*_self-reg_/*C*_const-rep_). Purple indicates that the self-regulating repressor results in lower cost. Circles mark points 1–3.

Here, the two optimized circuits have similar noise-free costs (within 2%), but reach them through different strategies. The constitutive-repressor circuit keeps a small repressor pool (*R* ≈ 2.6) and substantial basal resistance, so it is partly pre-induced before exposure. The self-regulated circuit maintains a much larger, self-repressed repressor pool (*R* ≈ 62) and keeps basal resistance close to zero until the drug arrives. Under these circumstances, repressor noise explains the weakness of constitutive repressor expression. Before any drug is present, transient drops in the small repressor pool of the constitutive-repressor circuit release large bursts of resistance, which more than double the basal level of resistance at *σ*_R_ = 1. After exposure, the same fluctuations severely spread the timing of induction, and 7–20% of cells fail to recover growth altogether (**Fig. 5L**). In contrast, cells with the self-regulated architecture show no spurious induction before exposure, follow nearly identical trajectories after it, and all survive (**Fig. 5M**). Averaged over the population, the self-regulated architecture then becomes the preferred architecture at the highest drug concentrations (*D* ≥ 224 for *σ*_R_ ≥ 0.5), a region that extends down to lower drug concentrations at strong repressor noise (**Fig. 5N**).

This advantage follows from a known property of negative autoregulation, which buffers fluctuations in the concentration of the regulator itself^44–46^. A transient drop in repressor relieves self-repression and increases repressor synthesis, restoring the repressor pool before resistance is spuriously released. In a deterministic setting, self-regulation of the repressor carries only the cost of an additional regulatory element. Under noise, it protects against both unnecessary induction before exposure and failed induction after exposure. Notably, this benefit appears only near the limit of tolerable drug exposures, where small differences in the timing of induction determine whether a cell recovers or enters growth arrest^7,36^.

Noise in the expression of resistance has a different effect (**Fig. 6**). Constitutive expression of resistance is the most sensitive to this noise. Under drug exposure, expression of resistance decreases together with growth, and cells that fluctuate to low levels of resistance enter growth arrest (**Fig. 6A, E, I**). As a result, a large fraction of cells with constitutive resistance collapse already at intermediate drug concentrations, with survival dropping below 50% for ≳ 70 at *σ*_P_ = 1 (**Fig. 6P**). Both repressor-based architectures are considerably more robust, since induction raises expression of resistance well above the level needed to clear the drug (**Fig. 6Q, R**). Failure to recover growth only happens at very strong noise (*σ*_P_ ≳ 5) or, for the constitutive repressor, at the highest drug concentrations (*D* ≥ 224). Between the two regulated architectures, noise in resistance changes the preferred architecture only at the highest drug concentrations, where the self-regulated architecture is again favored (**Fig. 6S**). Thus, noise in the regulator, rather than in the resistance protein itself, is what makes self-regulation of the repressor advantageous over a wider range of conditions.

**Figure 6.**
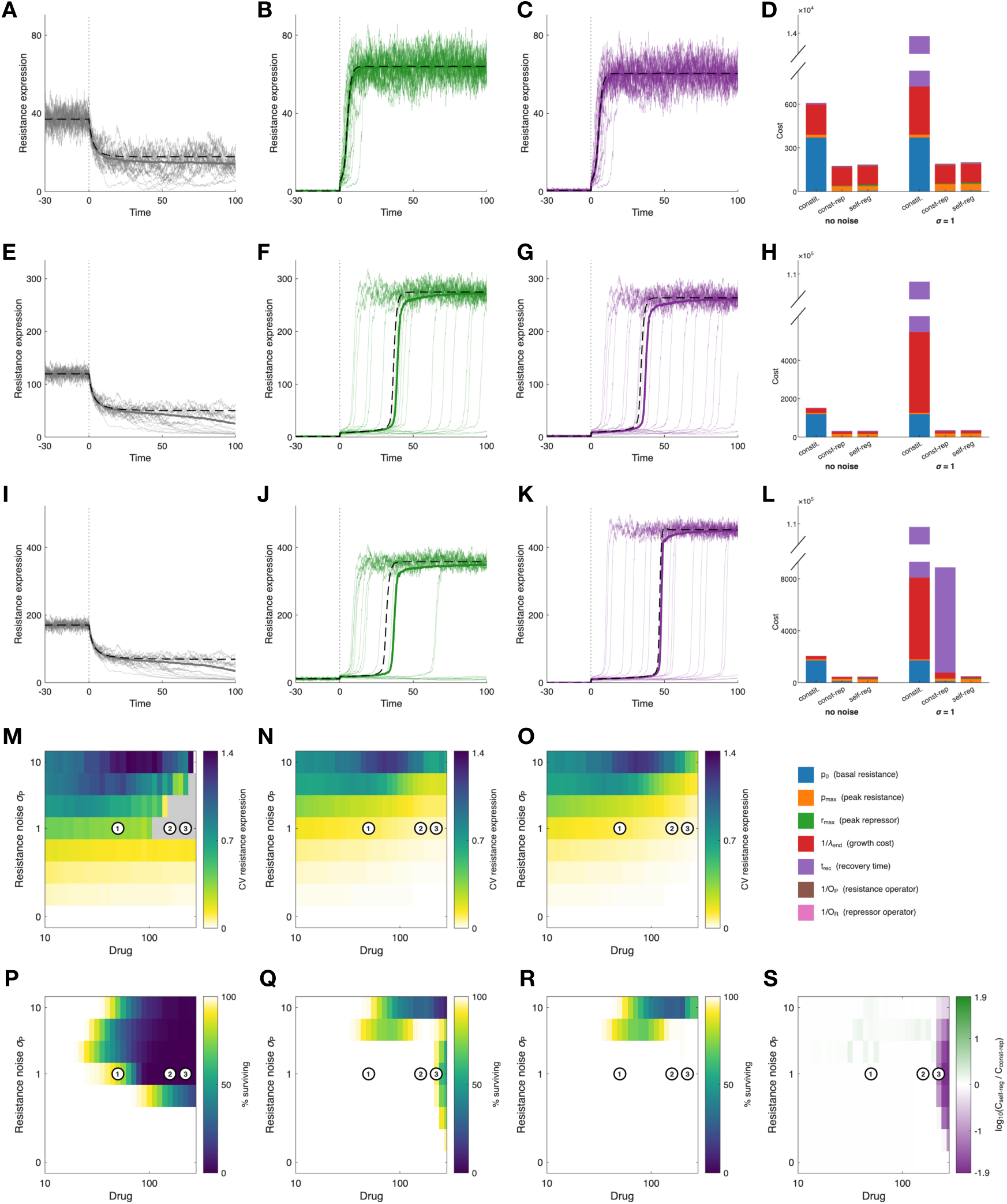
Noise in the expression of resistance penalizes constitutive resistance. Circuits were optimized as in Fig. 5, including constitutive resistance. (**A–L**) Responses to a drug exposure at time zero for *D* = 50 (**A–D**), *D* = 158 (**E–H**), and *D* = 224 (**I–L**), for constitutive resistance (**A, E, I**, gray), the constitutive repressor (**B, F, J**, green), and the self-regulating repressor (**C, G, K**, purple), with noise in the expression of resistance *σ*_P_ = 1, shown as in Fig. 5. (**D, H, L**) Breakdown of the cost without noise and with *σ*_P_ = 1, averaged over 1000 cells, including cells that collapse. (**M–O**) Cell-to-cell coefficient of variation of resistance at the end of exposure among surviving cells, for constitutive resistance (**M**), constitutive repressor (**N**), and self-regulating repressor (**O**); gray indicates no surviving cells. (**P–R**) Percentage of surviving cells. (**S**) Preferred architecture between the self-regulating and constitutive repressors, measured by log_10_(*C*_self-reg_/*C*_const-rep_). Constitutive resistance results in the highest cost at all drug concentrations and noise levels shown.

## Discussion

The regulatory strategy of an antibiotic response should be understood as an adaptation to the demands imposed by the environment. A resistance protein that protects the cell from the drug is also costly, and regulation can resolve this conflict. However, regulation is not free: repression delays the response, impairing growth, and imposes additional costs of its own. Our results show that changing the relative importance of these costs is sufficient to shift the optimal response between distinct regulatory architectures. Thus, selection acts not simply on the resistance protein itself, but on the full time course of the response.

The three architectures analyzed here can therefore be viewed as solutions to distinct dynamical problems, rather than as a simple progression in regulatory complexity. Constitutive expression of resistance keeps protection available at all times, and is favored when the cost of delayed activation outweighs the burden of pre-existing expression. Repression by a constitutive repressor becomes advantageous when basal expression of resistance is especially costly and exposures are sufficiently rare that the benefit of keeping resistance off exceeds the cost of slower activation. Self-regulation of the repressor adds further flexibility, but this flexibility is not required in most regimes. Its value lies in allowing the cell to adjust not only the level of protection, but also the way protection is deployed over the course of the response.

Experimental evolution under controlled drug dynamics has shown that the regimen of drug administration shapes the evolutionary paths to resistance^47–49^. Our predictions are supported by our recent experimental evolution of *E. coli* populations carrying the tetracycline resistance *tet* operon under contrasting regimens of drug administration^13^. Under gradually increasing tetracycline concentrations, populations preserved the native regulated response and instead acquired mutations in the efflux pump AcrB that increase its affinity for tetracycline. In contrast, under periodic sudden exposures to large drug doses, all populations acquired transposon insertions in repressor *tetR*, resulting in constitutive expression of efflux pump TetA. This contrast supports the prediction that, when the dynamics of the environment compress the time available for induction, the cost of delayed activation outweighs the benefit of keeping resistance off between exposures, shifting the optimum towards constitutive protection. More broadly, it shows that the dynamics of the environment can alter not only the optimal parameters of a response, but the optimal regulatory architecture itself, and may help explain why constitutive expression of resistance is frequently observed among clinical isolates^13,15^.

The self-regulating architecture shows that additional regulatory elements are used selectively. Repression of resistance is optimal over most conditions, but self-regulation of the repressor is recruited only at high drug concentrations and high costs of peak resistance. In this regime, self-regulation allows the cell to keep basal resistance close to zero while maintaining a large repressor pool, at the cost of a delayed induction followed by an overshoot in resistance. Therefore, self-regulation is not a requirement for efficient control, but a refinement that becomes valuable only in the most demanding regimes. However, this small advantage becomes decisive once noise in gene expression is taken into account.

Our stochastic simulations show that noise in gene expression further shapes the choice of architecture. Away from the limit of tolerable drug exposures, noise changes costs only modestly, and the deterministic optima remain good predictors of the preferred architecture. Near that limit, where the timing of induction determines whether a cell recovers or enters growth arrest^7,8^, noise in the repressor reshapes the choice of regulatory architecture. A constitutive repressor maintained at low levels is prone to spontaneous de-repression, which results in unnecessary induction of resistance before exposure and, after exposure, a spread of induction times in which some cells fail to recover. Self-regulation of the repressor buffers these fluctuations^44–46^, so that an architecture with a small cost penalty in the deterministic setting becomes the preferred one once noise is taken into account. This suggests a role for the strong self-repression observed in stress responses such as the *tet* and *mar* systems^37,50^, in addition to its established effects on the speed of the response^51–53^. By contrast, noise in the expression of resistance mainly penalizes constitutive expression, whose levels decrease together with growth during drug exposure, and has little effect on the choice between the two repressor-based architectures.

These design principles are consistent with the diversity of regulatory strategies observed in natural systems^26^. Stress responses, where the intracellular signal directly threatens growth or survival, are often associated with strong repression and, in many cases, with strong self-repression of the regulator. The *tet* operon and the *marRAB* system are examples where the response is tightly coupled to the harmful signal itself, and the timing of the response is a dominant design constraint^29,50,54,55^. The SOS response illustrates a related logic: when persistent damage makes constant readiness advantageous, self-repression can be weakened or lost altogether in some lineages^56,57^. Metabolic responses show a broader range of strategies. Responses such as IclR regulation of *aceBAK* and MtlR regulation of *mtlADR* resemble stress responses when rapid processing becomes important or when alternative resources are scarce^58–61^, while the *lac* operon is regulated by a nearly constitutively expressed repressor, in a regime where controlling expression levels is more important than rapidly processing the signal^62,63^. Galactose utilization provides a mixed example, where GalS and GalR regulate complementary transport and utilization programs under low and high galactose concentrations, respectively^64,65^. The same logic may also explain shifts in the architecture of antibiotic resistance mechanisms, such as the contrast between low constitutive expression of AmpC β-lactamase in some organisms and strongly regulated expression in others^66–68^. Together, these examples support the view that the same response motif can be embedded in distinct regulatory architectures, because different environments impose selective pressure on different phases of the response^69,70^.

Our model is most directly applicable to responses where a harmful intracellular drug is sensed by a transcription factor that regulates a resistance protein acting directly on that drug, and where sudden exposure drives the response out of equilibrium. Caution is required when extending the model to responses dominated by active import or by changes in membrane permeability^71,72^, transcriptional activation rather than repression, or complex downstream cascades. The present formulation also relies on simplifying assumptions, including rapid binding equilibrium, a reduced description of growth coupling, and a fixed Hill coefficient. Our analysis of noise treats fluctuations phenomenologically, as noise in synthesis scaled by a single amplitude factor, without explicitly resolving transcriptional bursting, partitioning at cell division, or extrinsic noise shared between genes^39,40^, and noise was evaluated on circuits optimized deterministically. Optimizing regulation directly against the population-averaged stochastic cost could shift the optima further, and coupling this framework to population dynamics would be necessary to translate survival into fitness. Nonetheless, these simplifications do not alter the main conclusion: regulatory architecture is an evolvable solution to a dynamical design problem, and different environments can favor loss, retention, or elaboration of regulation, according to which burdens dominate the response.

Describing the cell’s environment as a series of costs associated with the response, our framework captures the response as a temporally extended process rather than a sequence of steady states. The metrics that enter it include both steady-state measures (basal and final levels, and operator strength) and kinetic measures (transient peaks of resistance and repressor, and recovery time). We assumed that the cost grows linearly with each metric, which is sufficient to recover the qualitative trade-offs discussed above and is the simplest assumption consistent with our approach^26^. Nonlinear cost functions, such as saturating penalties or steeper penalties for growth arrest, would shift quantitative details such as the precise location of optima, but would not alter the conclusion that distinct selective regimes favor distinct architectures.

Our optimization identifies the optimal architecture and parameters in each environment, but evolution will reach these optima only along available mutational paths. The cost landscapes we examined are largely well-behaved, but contain extended, nearly flat regions where many parameter combinations result in similar costs, so the observed regulatory parameters may reflect drift or historical contingency as much as selection. Transitions between architectures, such as loss of repression or acquisition of self-regulation, depend on the available mutational mechanisms. Since regulatory mutations are typically more accessible than mutations in coding regions^73,74^, the architectures favored in dynamic environments may be not only the fittest, but also among the most readily reached.

Taken together, our results frame regulatory architecture as an evolvable adaptation to the dynamical demands of the environment. The same motif of a repressor-regulated resistance gene can be embedded in qualitatively different architectures (constitutive expression, repression by a constitutive repressor, or repression by a self-regulated repressor), and our framework identifies the conditions under which each is preferred. Including noise in this framework shows that the preferred architecture depends not only on the dynamics of the environment, but also on the fluctuations the response must tolerate: noise in the repressor can make self-regulation favorable in regimes where deterministic optimization would discard it, so that regulatory elements that appear dispensable on average may be maintained because they make the response reliable in individual cells. By treating the response as a dynamical process and the cost function as the integrator of competing selective pressures, this approach provides a basis to predict which regulatory strategies should arise in each environment, and may be useful both to interpret the diversity of natural regulatory systems and to guide the design of synthetic circuits^75,76^.

## Materials and methods

### Model

We model the concentrations of intracellular drug *d*, resistance protein *p*, and repressor *r*, in nondimensional form, as

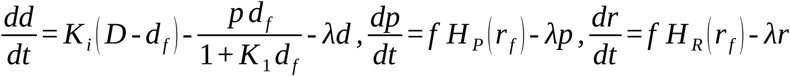

where *D* is the extracellular drug concentration, which increases from zero to a fixed value at time *t* = 0. Free drug *d*_f_ and free repressor *r*_f_ = *r*/(1 + *K*_2_*d*_f_) are calculated from the binding equilibrium between drug and repressor, with *d* = *d*_f_ + (*r* – *r*_f_)/*K*_2_. Growth is described by the reduced proteome partition model^8,32^, with *λ*(*d*_f_) = (1 + *K*_x_)/[1 + *K*_x_(1 + *d*_f_)] and *f* = *λ*^2^. Regulation of resistance and repressor synthesis is given by

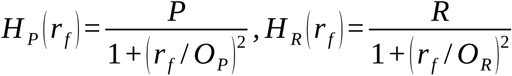

for the self-regulating repressor. For the constitutive repressor, *H*_R_ = *R*, and for constitutive resistance, *r* = 0 and *H*_P_ = *P*. Time is measured in units of 1/*λ*_0_, where *λ*_0_ is the drug-free growth rate, drug concentrations in units of the drug concentration for half-inhibition of translation, and repressor concentrations in units of the drug–repressor dissociation constant. We used *K*_i_ = 1, *K*_1_ = 0.1, *K*_2_ = 1000, and *K*_x_ = 0.47, based on the tetracycline response in *E. coli*^7,8^. Simulations start from the drug-free steady state and were integrated with ode113 in MATLAB until the system reached a stable steady state. The full dimensional model, the rescaling, and all parameter values are given in **Supplementary Text S1** and **Tables S1–S3**.

### Cost function

For each simulated response, we calculated the basal levels of resistance and repressor before exposure (*p*_0_, *r*_0_), their maximal levels during the response (*p*_max_, *r*_max_), the final growth rate *λ*_end_, and the recovery time *t*_rec_, defined as the time from the growth minimum until growth exceeds 80% of *λ*_end_. The cost of the response is

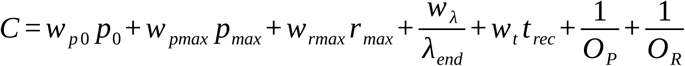

where operator penalties apply only to the operators present in each architecture. For constitutive resistance (**Fig. 2**), *C* = *w*_p0_*p*_0_ + 100/*λ*_end_. For the constitutive repressor (**Fig. 3**), *w*_pmax_ = *w*_rmax_ = *w*_t_ = 1 and *w*_λ_ = 100, and *w*_p0_ was varied. For the self-regulating repressor (**Fig. 4**), *w*_p0_ = 10, *w*_rmax_ = *w*_t_ = 1, *w*_λ_ = 100, and *w*_pmax_ was varied (**Table S5**). Responses with *λ*_end_ < 0.2, or that did not reach a steady state within *t* = 100, were considered unviable and assigned infinite cost (*λ*_end_ < 0.02 for constitutive resistance).

### Optimization

Drug concentrations were sampled on a logarithmic grid *D*_i_ = 10^5i/100^ (*i* = 1, …, 49; 1.1 ≤ *D* ≤ 282), and weights on a logarithmic grid *w*_j_ = 10^−1+4*j*/100^ (*j* = 1, …, 70; 0.11 ≤ *w* ≤ 63). At each grid point, regulatory parameters were optimized with a particle swarm algorithm^77^ (MATLAB particleswarm, 300 particles), followed by local refinement with a pattern search^78^ (patternsearch), within the bounds listed in **Table S5**. Points where the optimization converged to local minima, identified as discontinuities in the optimal parameters, were re-optimized starting from the optima of neighboring grid points. We verified the optima with an independent evaluator that only accepts stable steady states as the end of a response (**Supplementary Text S3**).

### Stochastic simulations

Noise in the synthesis of repressor or resistance was modeled with chemical Langevin equations^43^, adding a term *σ*(*f H*)^1/2^*ξ*(*t*) to the corresponding equation, where *ξ* is Gaussian white noise and *σ* = *σ*_R_ or *σ*_P_ sets the relative noise amplitude (*σ* = 0 and seven logarithmically spaced values between 0.1 and 10). Equations were integrated with a semi-implicit Euler–Maruyama scheme with time step 0.002, with noise starting 50 time units before exposure. Each condition was simulated for 1000 cells, using the same noise realizations for all architectures. Exposure lasted 300 time units for repressor noise and 1000 time units for resistance noise, to capture slow recoveries. Cells with mean growth below 0.2 over the last 10 time units were considered not surviving, and assigned a recovery time of 10^5^. Circuits were optimized under the cost of **Fig. 4** with *w*_pmax_ = 0.52, with the self-regulated circuit constrained to *r*_0_/*R* = 0.2 (**Supplementary Text S4**, **Table S6**).

## Data Availability

All code used to generate the figures in this article is available at our GitHub page: https://github.com/schultz-lab/regulatory-architectures. Further inquiries can be directed to the corresponding author.

## Author Contributions

H.G. and D.S. designed the study and developed the model. H.G. performed the optimizations and stochastic simulations. H.G. and D.S. analyzed the results and wrote the manuscript.

## Acknowledgements

We thank Philip Bittihn and Lev Tsimring for valuable discussions. We also thank Mirjana Stevanovic and João Pedro T. Carvalho for their help developing the model, as well as John Crow for his experimental work that supports this work.

