## Supplemental Information for "Gene regulatory architecture in antibiotic resistance is optimized for specific dynamic regimens"

### Supplementary Information

**Gene regulatory architecture in antibiotic resistance is optimized for specific dynamic regimens**
**Hao Geng[1] and Daniel Schultz[1]***
[1] Department of Microbiology & Immunology – Geisel School of Medicine at Dartmouth. Hanover, NH 03755, USA.

**Contents**

Supplementary Text S1. Model of the antibiotic response
Supplementary Text S2. Response metrics and cost function
Supplementary Text S3. Optimization of regulatory parameters
Supplementary Text S4. Stochastic simulations
Tables S1–S6
Supplementary References

#### Supplementary Text S1. Model of the antibiotic response

##### Dimensional model

We model the response by three differential equations describing the concentrations of total intracellular drug $D$, resistance protein $P$, and total repressor $R$[1,2]:

$$\frac{dD}{dt} = K_i\left(D_{out}(t) - D_f\right) - \frac{K_P\, P\, D_f}{k_P + D_f} - \lambda D$$

$$\frac{dP}{dt} = f\, H_P\left(R_f\right) - \lambda P$$

$$\frac{dR}{dt} = f\, H_R\left(R_f\right) - \lambda R$$

The drug enters the cell with rate constant $K_i$, proportionally to the difference between the extracellular concentration $D_{out}(t)$ and the free intracellular concentration $D_f$. It is processed by the resistance protein following Michaelis–Menten kinetics, with maximal processing rate $K_P$ and Michaelis constant $k_P$, and all components are diluted by cell growth at rate $\lambda$. Binding and unbinding of the drug to the repressor are much faster than synthesis, transport, and growth, so we consider the free forms of drug and repressor ($D_f$, $R_f$) to be in equilibrium with the bound complex $[RD]$, with dissociation constant $K_d$:

$$R_f\, D_f = K_d\,[RD], \qquad D = D_f + [RD], \qquad R = R_f + [RD]$$

Since the drug–repressor complex is inactive, only free repressor regulates transcription. Synthesis of resistance and repressor is regulated by free repressor according to Hill functions, with Hill coefficient $n = 2$ to reflect the binding of repressor dimers:

$$H_P\left(R_f\right) = \frac{\alpha_P}{1 + \left(R_f/R_p\right)^2}, \qquad H_R\left(R_f\right) = \frac{\alpha_R}{1 + \left(R_f/R_r\right)^2}$$

where $\alpha_P$ and $\alpha_R$ are the maximal synthesis rates (promoter strengths) and $R_p$ and $R_r$ are the concentrations of free repressor resulting in half-maximal repression of each promoter (operator thresholds). For the constitutive repressor, $H_R = \alpha_R$, and for constitutive resistance, there is no repressor and $H_P = \alpha_P$.

##### Effects of the drug on growth and gene expression

We describe the effects of the drug on growth and gene expression with the proteome partition framework of Scott *et al.*[3,4], in the reduced form used in our previous work[2]. Growth is determined by the translational capacity $\kappa_t$ and the nutritional capacity $\kappa_n$ of the cell, with $\lambda \propto \kappa_t\kappa_n/(\kappa_t + \kappa_n)$. We assume that the nutritional capacity remains at its drug-free value, while the translational capacity is reduced by free intracellular drug, $\kappa_t = \kappa_t^0/(1 + D_f/K_r)$, where $K_r$ is the drug concentration for half-inhibition of translation. The growth rate is then

$$\frac{\lambda}{\lambda_0} = \frac{1 + K_x}{1 + K_x\left(1 + D_f/K_r\right)}, \qquad K_x = \frac{\kappa_n}{\kappa_t^0}$$

where $\lambda_0$ is the drug-free growth rate. Resistance and repressor are assumed to belong to the proteome sector whose expression decreases with growth and translational inhibition, so their synthesis is modulated by the factor $f = (\lambda/\lambda_0)^2$, which accounts for both the decrease in growth and the reallocation of the proteome towards ribosomal proteins[2,5]. All parameters are listed in **Table S1**, and were chosen based on the tetracycline response in *E. coli* growing in M63 medium[1,2,6].

##### Nondimensional model

We rescale time by the drug-free growth rate, drug concentrations by $K_r$, repressor concentrations by $K_d$, and resistance by its drug-processing capacity (**Table S2**). Dropping primes, the model becomes

$$\frac{dd}{dt} = K_i\left(D - d_f\right) - \frac{p\, d_f}{1 + K_1 d_f} - \lambda d$$

$$\frac{dp}{dt} = f\,\frac{P}{1 + \left(r_f/O_P\right)^2} - \lambda p, \qquad \frac{dr}{dt} = f\,\frac{R}{1 + \left(r_f/O_R\right)^2} - \lambda r$$

with $\lambda = (1 + K_x)/[1 + K_x(1 + d_f)]$, $f = \lambda^2$, and composite parameters $K_i$, $K_1$, $K_2$, and $K_x$ defined in **Table S3**. The binding equilibrium becomes $r_f = r/(1 + K_2 d_f)$, with $d = d_f + (r - r_f)/K_2$, which results in

$$d_f = \frac{K_2 d - r - 1 + \sqrt{(K_2 d - r - 1)^2 + 4K_2 d}}{2K_2}$$

In the implementation, this expression and the corresponding expression for $r_f$ are written in a numerically stable form to avoid cancellation errors. The regulation of the response is described by four nondimensional parameters: promoter strengths $P$ and $R$, and operator thresholds $O_P$ and $O_R$ (**Table S2**). The constitutive repressor has parameters $P$, $R$, and $O_P$, and constitutive resistance has only $P$.

##### Drug-free steady state and simulation of the response

In the absence of drug, $\lambda = f = 1$ and $r_f = r$. The basal level of repressor is $r_0 = R$ for the constitutive repressor, and the solution of $r_0 = R/[1 + (r_0/O_R)^2]$ for the self-regulating repressor. The basal level of resistance is $p_0 = P/[1 + (r_0/O_P)^2]$ (or $p_0 = P$ for constitutive resistance). Starting from this steady state, the extracellular drug concentration is raised from zero to $D$ at $t = 0$, and the equations are integrated with ode113 (MATLAB; relative and absolute tolerances $10^{-7}$ and $10^{-9}$). The fixed points of the system in the presence of the drug are found by root finding from initial guesses $d = D$ and $d = 0$, which correspond to the arrested and recovered states when the system is bistable[2,7]. Integration stops when the state reaches a fixed point within 10% (relative norm), and the final levels of all components are taken at that fixed point.

#### Supplementary Text S2. Response metrics and cost function

From each simulated response we extract the metrics listed in **Table S4**. The cost of the response is a weighted sum of these metrics,

$$C = w_{p0}\,p_0 + w_{r0}\,r_0 + w_{pmax}\,p_{max} + w_{rmax}\,r_{max} + w_{pend}\,p_{end} + w_{rend}\,r_{end} + \frac{w_\lambda}{\lambda_{end}} + w_t\,t_{rec} + \frac{1}{O_P} + \frac{1}{O_R}$$

where operator penalties apply only to the operators present in each architecture. The weights used for each figure are listed in **Table S5**. In all scenarios considered here, $w_{r0} = w_{pend} = w_{rend} = 0$. We assume that the cost increases linearly with each metric, which is the simplest assumption that captures the trade-offs between the different burdens of the response[1]. The growth cost enters as $1/\lambda_{end}$, so that responses that maintain growth have lower cost. The recovery time $t_{rec}$ is measured from the minimum in growth rate after exposure until growth first exceeds 80% of $\lambda_{end}$; if growth does not recover before the end of the simulation, $t_{rec}$ is the time at which the simulation ends. Responses whose final growth rate is below 0.2 (0.02 for constitutive resistance), or that do not reach a fixed point within $t = 100$, are considered unviable and assigned infinite cost. The maximal levels $p_{max}$ and $r_{max}$ are taken over the full response, including the approach to the final steady state. For the constitutive repressor, $r_{max}$ was taken as the steady-state level of repressor in the presence of the drug.

##### Cost decomposition for constitutive resistance (Fig. 2E)

For constitutive resistance, $p_0 = p_{max} = P$, and the cost reduces to $C = w_{p0}P + 100/\lambda_{end}(D, P)$. To illustrate the trade-off underlying the optimization, we fixed $D = 100$ and evaluated the two terms of the cost for levels of resistance between 0 and 150, for $w_{p0}$ = 10, 30, and 50. The cost of resistance increases linearly with $P$, while the growth cost decreases sharply once resistance is sufficient for the cell to escape the arrested state, resulting in two local minima of the total cost.

#### Supplementary Text S3. Optimization of regulatory parameters

For each architecture, the regulatory parameters were optimized on a two-dimensional grid of drug concentrations and weights. Drug concentrations were sampled as $D_i = 10^{5i/100}$, $i = 1, \ldots, 49$ ($1.12 \le D \le 282$), and weights as $w_j = 10^{-1+4j/100}$, $j = 1, \ldots, 70$ ($0.11 \le w \le 63.1$), resulting in 3430 grid points per architecture. The swept weight was $w_{p0}$ for constitutive resistance (**Fig. 2**) and for the constitutive repressor (**Fig. 3**), and $w_{pmax}$ for the self-regulating repressor (**Fig. 4**) (**Table S5**).

At each grid point, the cost was minimized with a particle swarm algorithm[8] (MATLAB particleswarm, 300 particles), followed by local refinement with a generalized pattern search[9] (MATLAB patternsearch, used as the hybrid function of particleswarm), within the bounds listed in **Table S5**. Parameters were optimized as $P$, $R$, $1/O_P$, and $1/O_R$. Removal of a regulatory element by the optimization corresponds to a parameter reaching its lower bound (for instance, $1/O_R \to 10^{-4}$ makes the self-regulating repressor equivalent to a constitutive repressor). Because the cost landscape contains discontinuities caused by the bistability between recovered and arrested states (**Fig. 2E**), the optimization occasionally converged to local minima, identified as isolated discontinuities in the maps of optimal parameters. These points were re-optimized within narrower bounds and starting from the optima of neighboring grid points, and the lower of the two costs was kept.

##### Verification with a horizon-free evaluator

The simulations used for optimization stop at the first approach to any fixed point of the system. At high drug concentrations, where the system is bistable, a trajectory can pass slowly near the unstable fixed point (saddle) before collapsing to the arrested state, and would be scored as a recovery. To verify the optima, we re-evaluated all optimized responses with an independent evaluator that integrates the equations until the state is stationary, checks the stability of the fixed point reached through its Jacobian, and only ends the response at a stable fixed point. All cost terms and the requirement that the response reaches its fixed point by $t$ = 100 were kept unchanged. This evaluator reproduces the original costs to within 0.07% wherever both agree on the outcome of the response. For constitutive resistance and the constitutive repressor (**Figs. 2** and **3**), no optima were affected. For the self-regulating repressor (**Fig. 4**), 262 of 3430 points were affected: 24 points where a passage near the saddle was scored as a recovery ($71 \leq D \leq 89$), and 238 points that reached the stable fixed point slightly after the $t$ = 100 limit ($50 \leq D \leq 282$, $w_{pmax} \geq 6.3$). Re-optimizing these points changed their costs by a median of –0.03% (maximum +8.5%), so **Fig. 4** was kept unchanged. However, the optimal parameters changed by more than two-fold at 203 of these points, reflecting the nearly flat cost landscape in the region where self-regulation is used.

#### Supplementary Text S4. Stochastic simulations

##### Noise model

To determine how noise in gene expression affects the cost of each architecture, we added noise to the synthesis of either the repressor or the resistance protein, using chemical Langevin equations[10,11]. For repressor noise, the repressor equation becomes

$$dr = \left[f\, H_R(r_f) - \lambda r\right]dt + \sigma_R \sqrt{f\, H_R(r_f)}\; dW_r$$

and analogously for resistance noise, with amplitude $\sigma_P$, where $W$ is a Wiener process. The amplitude of fluctuations therefore scales with the square root of the synthesis rate, as expected for Poisson-distributed synthesis events, and the relative amplitude $\sigma$ sets the effective size of synthesis events in the scaled units of the model. We considered $\sigma = 0$ and seven logarithmically spaced values between 0.1 and 10.

##### Numerical integration

Equations were integrated with a semi-implicit Euler–Maruyama scheme with time step $\Delta t = 0.002$. Synthesis and noise were treated explicitly, and dilution and drug processing implicitly:

$$x_{n+1} = \frac{x_n + fH\,\Delta t + \sigma\sqrt{fH}\,\sqrt{\Delta t}\;\xi_n}{1 + \lambda\,\Delta t}, \qquad d_{n+1} = \frac{d_n + K_i D\,\Delta t}{1 + k_{eff}\,\Delta t}$$

where $x$ is the noisy species, $\xi_n$ are independent standard normal variables, and $k_{eff} = [K_i d_f + p\, d_f/(1 + K_1 d_f) + \lambda d]/d$. Concentrations were truncated at zero. Larger time steps ($\Delta t$ = 0.005 and 0.01) changed the outcome of some responses near the threshold for collapse at high drug concentrations, while $\Delta t$ = 0.002 agreed with deterministic integration with ode15s.

#### Simulation protocol and metrics

Each simulation started at the drug-free steady state and ran for 50 time units with noise and no drug, so that the state of each cell at the time of exposure includes any spontaneous de-repression of resistance. The drug was then added at concentration $D$, and the response was simulated for 300 time units (repressor noise) or 1000 time units (resistance noise, where slow recoveries are more frequent). For each cell, $p_0$ and $r_0$ were taken at the time of exposure, $p_{max}$ and $r_{max}$ as the maxima during exposure, and $\lambda_{end}$ as the mean growth rate over the last 10 time units. The recovery time was calculated as in the deterministic case, within the period before the state (smoothed over 2 time units) first reaches within 10% of its final mean state. Cells with $\lambda_{end} < 0.2$ were considered not surviving and assigned $t_{rec} = 10^5$, corresponding to the infinite cost of unviable responses in the deterministic optimization. The cost of each condition is the mean cost over 1000 cells, including cells that do not survive. All architectures were simulated with the same sequence of random numbers (common random numbers), so that differences between architectures are not obscured by sampling noise. We also recorded the fraction of surviving cells, the induction time of each surviving cell (the time at which resistance first reaches halfway between its level at exposure and its final level), and the cell-to-cell coefficient of variation of resistance at the end of exposure among surviving cells.

#### Optimized circuits

Circuits for the stochastic simulations were optimized under the cost of **Fig. 4** ($w_{p0} = 10$, $w_{rmax} = w_t = 1$, $w_\lambda = 100$) with $w_{pmax} = 0.52$ (grid row $j = 18$), using the horizon-free evaluator (**Supplementary Text S3**). The constitutive repressor was optimized over $P$, $R$, and $1/O_P$, and constitutive resistance over $P$. Since optimization of the self-regulating repressor converges to a constitutive repressor at this weight, we constrained the strength of self-repression to $r_0/R = \varepsilon = 0.2$, which, from the steady-state condition, fixes $1/O_R = k/R$ with $k = (1/\varepsilon - 1)^{1/2}/\varepsilon = 10$, and optimized $P$, $R$, and $1/O_P$. The value $\varepsilon = 0.2$ is the median of $r_0/R$ over the region of **Fig. 4** where self-regulation is used ($r_0/R < 0.85$; median 0.21, mean 0.28). Optimization used particle swarm (40 particles, 40 iterations) seeded with the constitutive-repressor optimum and the optima at neighboring drug concentrations, followed by pattern search, and a final pass in which each point was re-optimized starting from the optima of its neighbors. The optimized circuits at the three points shown in **Figs. 5** and **6** are listed in **Table S6**. In the absence of noise, the self-regulated circuit costs 6.3% more than the constitutive repressor at low drug concentrations, decreasing to 1.0% at $D = 224$ and 0.1% at $D = 282$. At $D \geq 251$, the optimal promoter strength of resistance of the self-regulated circuit approaches the upper bound $P = 1000$, and the operator threshold $O_P$ of the self-regulated circuit is at its lower bound ($1/O_P = 5$) at all three points of **Table S6**.

#### Choice of weight

To choose the weight on maximal resistance for the stochastic analysis, we scouted ten weights between $w_{pmax} = 0.17$ and 43.7 with 200 cells per condition. Under repressor noise, the self-regulated circuit is cheaper than the constitutive repressor at high drug concentrations for $w_{pmax} \lesssim 0.5$, while for $w_{pmax} \geq 1.1$ both regulated architectures collapse at high drug concentrations and constitutive resistance becomes the cheapest. $w_{pmax} = 0.52$ is the weight where differences in both cost and survival between the two regulated architectures are apparent.

**Table S1. Parameters of the dimensional model.**

| Parameter | Value | Description |
|---|---|---|
| $K_i$ | 0.015 $min^{-1}$ | drug import rate |
| $K_P$ | 50 $min^{-1}$ | maximal drug-processing rate of the resistance protein |
| $k_P$ | 10 μM | Michaelis constant of drug processing |
| $K_d$ | 0.001 μM | drug–repressor dissociation constant |
| $K_r$ | 1 μM | drug concentration for half-inhibition of translation |
| $\kappa_t^0$ | 0.075 $min^{-1}$ | drug-free translational capacity |
| $\kappa_n$ | 0.035 $min^{-1}$ | nutritional capacity (M63 medium) |
| $\lambda_0$ | 0.015 $min^{-1}$ | drug-free growth rate (M63 medium) |
| $n$ | 2 | Hill coefficient of repression |
| $\alpha_P$, $\alpha_R$ | optimized | maximal synthesis rates of resistance and repressor (promoter strengths) |
| $R_p$, $R_r$ | optimized | free repressor for half-maximal repression of resistance and repressor promoters (operator thresholds) |

Parameter values follow our previous models of the tetracycline response in *E. coli*[1,2].

**Table S2. Nondimensional variables and regulatory parameters.**

| Quantity | Nondimensional form | Interpretation |
|---|---|---|
| time $t$ | $t' = \lambda_0 t$ | time in units of the inverse drug-free growth rate |
| intracellular drug $D$, $D_f$ | $d = D/K_r$, $d_f = D_f/K_r$ | drug relative to half-inhibition of translation |
| extracellular drug $D_{out}$ | $D' = D_{out}/K_r$ | drug concentration used in the figures |
| repressor $R$, $R_f$ | $r = R/K_d$, $r_f = R_f/K_d$ | repressor relative to drug–repressor dissociation constant |
| resistance $P$ | $p = K_P P/(k_P \lambda_0)$ | resistance relative to its drug-processing capacity |
| growth rate $\lambda$ | $\lambda' = \lambda/\lambda_0$ | growth relative to drug-free growth |
| promoter strength $\alpha_P$ | $P = K_P \alpha_P/(k_P \lambda_0^2)$ | maximal synthesis of resistance |
| promoter strength $\alpha_R$ | $R = \alpha_R/(K_d \lambda_0)$ | maximal synthesis of repressor |
| operator thresholds $R_p$, $R_r$ | $O_P = R_p/K_d$, $O_R = R_r/K_d$ | free repressor for half-maximal repression |

Primes are dropped in the main text and in Supplementary Text S1–S4.

**Table S3. Composite nondimensional parameters.**

| Parameter | Definition | Value | Interpretation |
|---|---|---|---|
| $K_i$ | $K_i/\lambda_0$ | 1 | drug import relative to growth |
| $K_1$ | $K_r/k_P$ | 0.1 | saturation of drug processing |
| $K_2$ | $K_r/K_d$ | 1000 | inactivation of repressor by the drug |
| $K_x$ | $\kappa_n/\kappa_t^0$ | 0.467 | coupling between translational capacity and growth |

**Table S4. Metrics composing the cost function.**

| Metric | Definition | Selective pressure represented |
|---|---|---|
| $p_0$ | basal resistance before exposure | cost of expressing resistance in the absence of drug; high weight favors repression by default |
| $r_0$ | basal repressor before exposure | cost of maintaining a large repressor pool |
| $p_{max}$ | maximal resistance during the response | cost of transient overexpression of resistance |
| $r_{max}$ | maximal repressor during the response | cost of transient overexpression of repressor |
| $p_{end}$, $r_{end}$ | final levels under sustained exposure | long-term cost of the response (not weighted in this study) |
| $1/\lambda_{end}$ | inverse of the final growth rate | growth inhibition under sustained exposure |
| $t_{rec}$ | time from the growth minimum until growth exceeds 80% of $\lambda_{end}$ | delay in growth recovery |
| $1/O_P$, $1/O_R$ | inverse operator thresholds | difficulty of evolving tight operators |

**Table S5. Cost function weights and optimization bounds.**

| Figure | Architecture | Fixed weights | Swept weight | Optimized parameters (bounds) |
|---|---|---|---|---|
| Fig. 2 | constitutive resistance | $w_\lambda = 100$ | $w_{p0}$ | $P$ (≤ 1000) |
| Fig. 3 | constitutive repressor | $w_{pmax} = w_{rmax} = w_t = 1$; $w_\lambda = 100$ | $w_{p0}$ | $P \in [10^{-4}, 1000]$<br>$R \in [10^{-4}, 1000]$<br>$1/O_P \in [10^{-4}, 5]$ |
| Fig. 4 | self-regulating repressor | $w_{p0} = 10$; $w_{rmax} = w_t = 1$; $w_\lambda = 100$ | $w_{pmax}$ | $P \in [10^{-4}, 1000]$<br>$R \in [10^{-4}, 500]$<br>$1/O_P$, $1/O_R \in [10^{-4}, 5]$ |
| Figs. 5, 6 | all three | as in Fig. 4 | $w_{pmax} = 0.52$ (fixed) | as in Figs. 2–4; self-regulating repressor with $r_0/R = 0.2$ |

Weights not listed are zero. All weights are swept on the grid $w_j = 10^{-1+4j/100}$, $j = 1, \ldots, 70$.

**Table S6. Optimized circuits used in the stochastic simulations (points 1–3 in Figs. 5 and 6).**

| $D$ | Architecture | $P$ | $R$ | $O_P$ | $O_R$ | Cost (no noise) |
|---|---|---|---|---|---|---|
| 50 | constitutive resistance | 37.0 | – | – | – | 609 |
| | constitutive repressor | 80.8 | 4.31 | 0.23 | – | 176 |
| | self-regulating repressor | 77.5 | 12.5 | 0.20 | 1.25 | 185 |
| 158 | constitutive resistance | 120 | – | – | – | 1520 |
| | constitutive repressor | 328 | 4.21 | 0.25 | – | 317 |
| | self-regulating repressor | 319 | 12.9 | 0.20 | 1.29 | 326 |
| 224 | constitutive resistance | 170 | – | – | – | 2061 |
| | constitutive repressor | 433 | 2.57 | 0.42 | – | 453 |
| | self-regulating repressor | 666 | 62.2 | 0.20 | 6.21 | 458 |

Circuits optimized under the cost of Fig. 4 with $w_{pmax} = 0.52$, using the horizon-free evaluator. The self-regulating repressor is constrained to $r_0/R = 0.2$. Costs from the deterministic optimization; the noise-free costs in Figs. 5 and 6, calculated with the stochastic simulator at $\sigma = 0$, differ by less than 1.5%.
